# Heartbeat related activity in the anterior thalamus differs between phasic and tonic REM sleep periods

**DOI:** 10.1101/2024.09.30.615986

**Authors:** Péter Simor, Róka Zita Lilla, Orsolya Szalárdy, Zsófia Jordán, László Halász, Loránd Erőss, Dániel Fabó, Róbert Bódizs

## Abstract

Rapid eye movement (REM) sleep is a fundamental sleep state associated with diverse functions from elemental physiological processes to higher order neurocognitive functions. A growing body of research indicates that REM sleep with eye movements (phasic REM) differs from REM periods without ocular activity (tonic) in terms of spontaneous and evoked neural responses. Studies using auditory stimulation consistently observed enhanced evoked responses in tonic versus phasic REM, indicating that the external processing is largely diminished when the eyes move during REM sleep. Whereas exteroceptive processing during sleep is widely studies, investigation on interoception (the processing of bodily signals) during sleep is scarce, and limited to scalp electroencephalographic recordings. Here we studied interoceptive processing in a group of epileptic patients (N = 11) by measuring their heartbeat-related neural activity in the anterior nucleus of the thalamus (ANT) during phasic and tonic REM sleep and resting wakefulness. Evoked potentials and beta–low gamma spectral power locked to the heartbeat were significantly different in phasic REM compared to tonic REM and wakefulness. Heartbeat-related neural signals exhibited pronounced inter-trial phase synchronization at lower (7–20 Hz) oscillatory activity in all vigilance states, but reduced gamma synchronization at later time points in phasic REM only. Tonic REM and wakefulness did not show significant differences in heartbeat-related activity in the ANT. Our findings indicate that heartbeat-related neural activity is detectable at the level of the ANT, showing distinct signatures of interoceptive processing in phasic REM compared to tonic REM and wakefulness.

## Introduction

Rapid eye movement (REM) is a fundamental sleep state that occupies around 20% of the total sleep time in human adults (Carskadon and Dement, 2005) and seems to play prominent roles in various biological functions including neurodevelopment, synaptogenesis, neural pruning, as well as cognition and affect (Blumberg et al., 2013; Datta, 2004; Li et al., 2017; Smith, 2004; van der Helm et al., 2011). Compared to non-REM sleep, REM sleep is characterized by mostly low-amplitude, high-frequency electroencephalographic (EEG) activity, alongside the transient appearance of slower frequency activities (such as bursts of 1–4 Hz delta waves and 4–7 Hz sawtooth waves). Additionally, irregularities in respiration and heartbeat are common during REM sleep (Blumberg et al., 2020). The combination of intense cortical and mental (dream) activity with muscular atonia makes this state appear peculiar, which is why it was also named paradoxical sleep (Jouvet, 1965). REM sleep features the alternation of periods with and without rapid ocular activity, often referred to as phasic and tonic REM states (Simor et al., 2020). Periods of rapid eye movements (phasic REM) are linked to neural activity originating from the pons (laterodorsal peduncolopintine tegmental nuclei) and propagating to the cortex (Fernández-Mendoza et al., 2009); however, phasic REM may also be initiated from top-down cortical projections extending to deeper regions (cortico-hypothalamic projections) (Hong et al., 2023).

Phasic and tonic REM periods alternate approximately every 60–70 seconds in humans (Bueno-Junior et al., 2023), and are proposed to represent two core aspects of sleep: environmental disconnection during phasic and external monitoring during tonic REM (Simor et al., 2020). Evidence supporting the distinct roles of phasic and tonic REM periods comes from studies aiming to probe cortical reactivity to external stimulation during REM sleep. These findings indicate that arousal and awakening thresholds are lower in tonic compared to phasic REM, the latter even reaching levels of the deepest stage of NREM sleep (Ermis et al., 2010). Moreover, acoustic stimuli during REM sleep appeared to elicit heightened cortical responses in tonic versus phasic REM sleep (REM), suggesting that sensory information arriving from the external environment is selectively suppressed when the eyes move during REM sleep (Koroma et al., 2020; Takahara et al., 2006). Attenuated reactivity to sensory stimulation during phasic REM was also corroborated in an fMRI study that examined whole-brain blood oxygen level dependent (BOLD) responses in REM sleep under acoustic stimulation (Wehrle et al., 2007). Whereas residual activity in auditory processing regions during tonic REM sleep were elicited to some extent and resembled those observed during resting wakefulness, in phasic REM periods no such responses were apparent. On the other hand, phasic REM featured the activity of a widespread thalamocortical network which appeared to be functionally isolated from external stimulation (Wehrle et al., 2007).

Reactivity to environmental stimuli during sleep is not limited to external sensory inputs. The perception of the internal state of the body through diverse (e.g., chemical, somatosensory, visceral or neural) ascending pathways, known as interoception, is critical to generate the cortical representation of the body, which in turn, functions as an interface between the brain and the body to adaptively regulate homeostatic functions, as well as to contribute to motivated behavior and affective states (Feldman et al., 2024; Seth, 2013). Although our sense of our body seems to be lost when we fall asleep, empirical studies indicate that interoceptive processing is maintained by the sleeping brain to some extent (Mazza et al., 2012), and seems to influence the regulation of sleep and arousal (Bastuji et al., 2008). Heartbeat-related cortical activity is one of the most studied measures of interoceptive processing (Coll et al., 2021). Cortical responses locked to the R-peak of the heartbeats vary in different states of vigilance and arousal, and also across phasic and tonic REM sleep (Immanuel et al., 2014, 2014; Simor et al., 2021a). More specifically, heartbeat-related potentials measured at the level of the scalp were not different in tonic REM and resting wakefulness, but differed in phasic REM periods at late (∼ 500 ms after the R-peak) components (Simor et al., 2021a).

Although heartbeat-related cortical activity is a widely used measure of interoceptive processing, only few studies assessed heartbeat-related activity beyond the level of the scalp (Park et al., 2018). Nevertheless, the processing and integration of bodily signals relies on extended subcortical pathways involving structures lying deep below the scalp such as the nucleus of the solitary tract, the parabrachial nucleus, the periaqueductal grey, to reach multimodal cortical sites such as the insula, and fronto-limbic regions through the thalamus (Berntson and Khalsa, 2021; Feldman et al., 2024; Park and Blanke, 2019).

Here, we studied heartbeat-related activity in REM sleep and resting wakefulness by assessing local field potentials of the anterior nucleus of the thalamus (ANT) in a group of epileptic patients undergoing surgery for deep brain stimulation. The ANT due to its anatomical connections is assumed to play a key role in the transmission of subcortical activations to cortical regions (Child and Benarroch, 2013; Gonzalo-Ruiz et al., 1995; Vertes et al., 2001). Moreover, as part of the fronto-limbic network, the ANT is intimately linked to brain regions (eg., amygdala, ventromedial prefrontal and anterior cingulate cortex) that orchestrate REM sleep (Hasegawa et al., 2022; Luppi et al., 2011; Miyauchi et al., 2009), and are also involved in the integration of interoceptive signals (Boccia et al., 2023; Feldman et al., 2024; Pollatos et al., 2007), making it a suitable diencephalic site below the cortex to study heartbeat-related activity in different vigilance states. In line with the notion of phasic REM as a state of environmental disconnection which also involves interoceptive processing, we expected distinct heartbeat-related neural responses in the ANT in phasic REM compared to tonic REM and wakefulness.

## Methods

### Participants

We examined the data from 12 epilepsy patients (mean age = 35.33 years, range: 17–64 years; 7 females) who were undergoing ANT deep brain stimulation (DBS) treatment at the National Institute of Clinical Neurosciences in Budapest, Hungary. Patients were selected from a larger pool (N = 20) based on the following inclusion criteria: one (seizure-and stimulation-free) recording night of co-registered ANT local field potentials (LFPs) and electrocardiography (ECG) was available, along with sufficient amounts (at least 5 minutes) of both phasic and tonic REM sleep segments in their nocturnal recordings, and similar amount of awake resting state recordings. One patient was excluded due to noisy ECG signal hindering the analyses of heartbeat related activity. The research protocol was approved by the local ethical committee of the National Institute of Clinical Neurosciences, and all participants provided informed consent. Clinical and demographic data are reported in **Table 1**.

**Table1.**
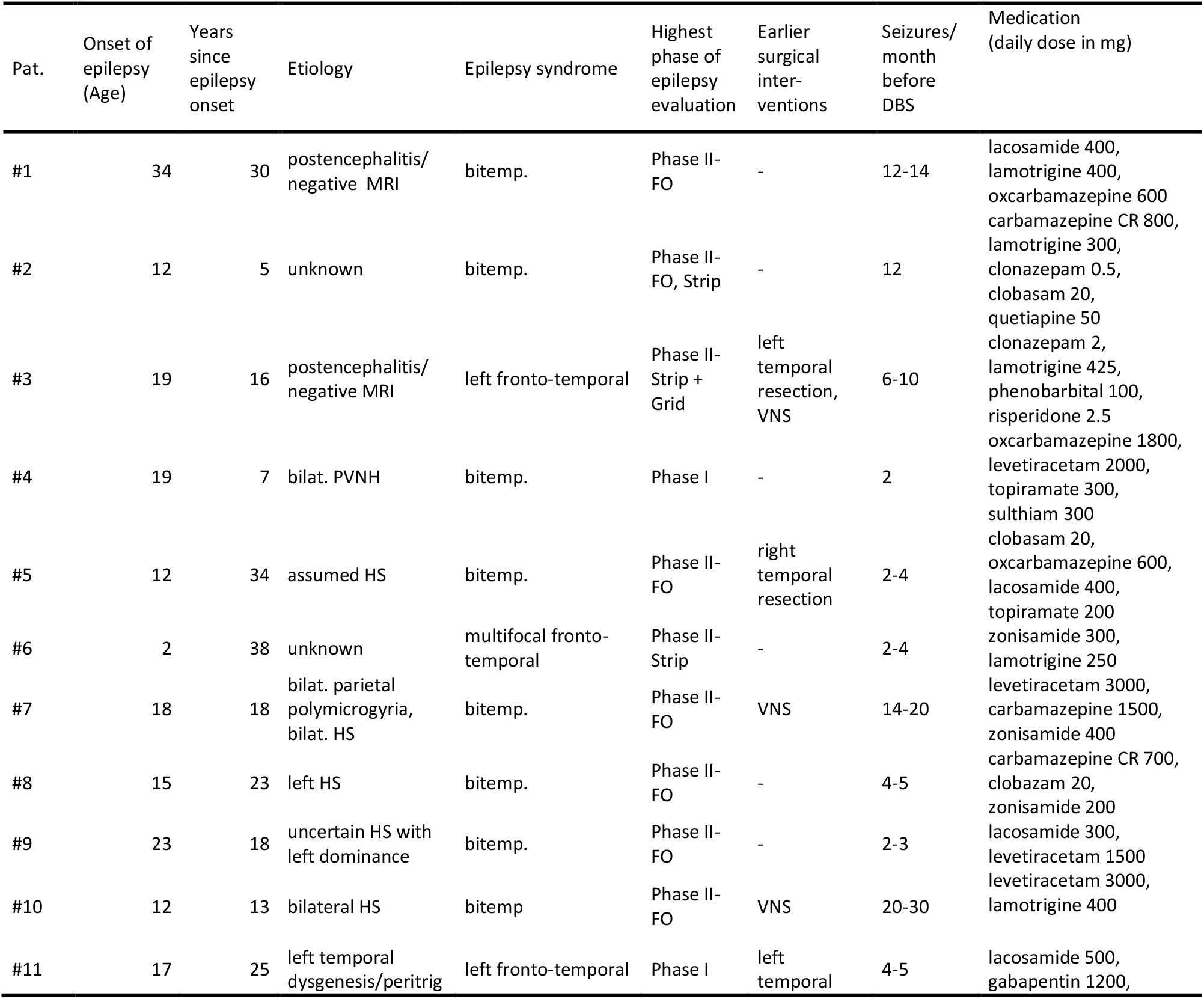
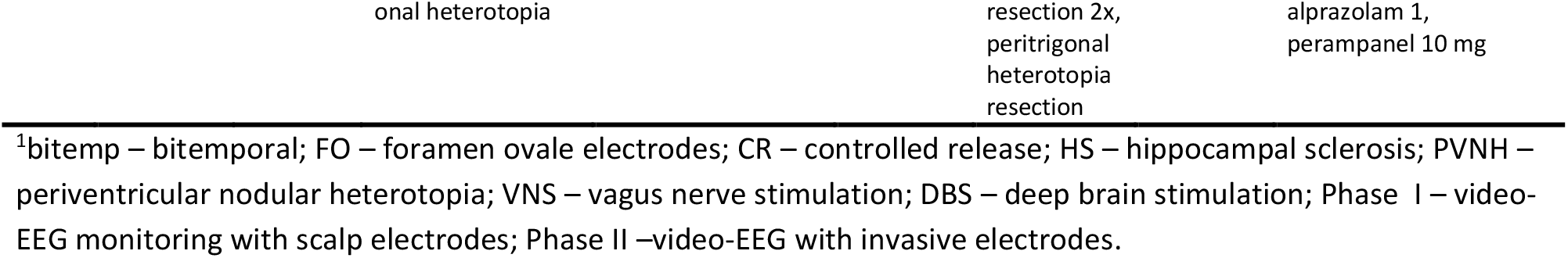
Clinical data of the subjects involved in the study.

### Surgical Procedures

Medtronic DBS electrodes were stereotactically implanted bilaterally in the anterior thalamus under general anesthesia to mitigate seizures in pharmacoresistant, surgically untreatable epilepsy patients. In addition to mostly frontal transventricular trajectories (nine patients), extraventricular trajectories, and posterior parietal extraventricular approaches were employed based on the decisions of the clinical-neurosurgical team.

### Localization of ANT channels

Preoperative MRI and postoperative CT images were co-registered using tools from the FMRIB Software Library (FSL, Oxford, FLIRT, linear registration, 6 degrees of freedom) to ensure precise localization of thalamic contacts. A threshold was applied to the co-registered CT scans to achieve the desired density for proper lead identification, thus excluding the surrounding brain tissue. The coordinates of the lead’s most distal point were identified, and a more proximal point was selected along the line of contacts to mathematically reconstruct the centre point coordinates of each contact using Euclidean distance in three-dimensional space. These points, superimposed on the T1 MRI image, provided a guideline for contact localization by examining their position relative to the anatomical boundaries of the ANT (see (Simor et al., 2021b) for more details). The anatomical positions of the contacts were double-checked using the mamillothalamic tract as a guide to localize the ANT. When both methods produced convergent results, the ANT contacts were considered suitable for further analyses.

### Electrophysiological recordings

Externalized DBS electrodes were connected to an EEG headbox/amplifier. Electrophysiological and video recordings were performed by using an SD-LTM 64 Express EEG/polygraphic recording system and the System Plus Evolution software (Micromed). Signals were recorded at 8192Hz/channel effective sampling rate with 22-bit precision and hardware input filters set at 0.02 (high pass: 40 dB/decade) and 450 Hz (low pass: 40 dB/decade). Data were decimated by a factor of four by the firmware resulting in stored time series digitized at 2048Hz/channel with the exception of one patient where it was set to 1024Hz/channel due to technical issues. LFPs of the ANT were assessed by bilateral (L, left; R, right) quadripolar electrodes (LTh0, LTh1, LTh2, LTh3, RTh8, RTh9, RTh10, RTh11) according to a bipolar reference scheme: LTh0-LTh1, LTh1-LTh2, LTh2-LTh3, LTh0-LTh3, RTh8-RTh9, RTh9-RTh10, RTh10-RTh11, RTh8-RTh11. LTh0-LTh3 and RTh8-RTh11 were the distant reference derivations on the left and the right side of the ANT, respectively. Scalp EEG allowing sleep staging was recorded following standard protocols (Keil et al., 2014). Scalp EEG was recorded at electrode placements according to the 10–20 system (Fp1, Fp2, Fpz, F3, F4, F7, F8, C3, C4, T3, T4, T5, T6, P3, P4, O1, O2, Oz) extended with the inferior temporal chain (F9, F10, T9, T10, P9, P10) and additional two anterior zygomatic electrodes (ZA1, ZA2; (Manzano et al., 1986)). EEG signals were re-referenced offline to the mathematically linked T9 and T10 points [(T9 + T10)/2]. Ocular movements facilitating the identification of phasic and tonic segments were assessed by the ZA1-ZA2 bipolar record created by off-line re-referencing. Submental electromyograms (EMGs) were recorded by bipolarly referenced electrodes placed on the chin. ECG activity was recorded by an electrode placed at the midclavicular line on the left and the fifth intercostal space referred to the CP1 location on the scalp.

### Data preprocessing

Data and statistical analyses were performed in MATLAB, using custom-based scripts and functions from the Fieldtrip toolbox (Oostenveld et al., 2011) and in JASP (0.18.3) [Team 2020]. Acquisition of stimulation-free nocturnal recordings after the surgical implantation of thalamic contacts was described in detail previously (Simor et al., 2021b; Szalárdy et al., 2024). In sum, continuous EEG and LFP recordings were segmented into 90-minute chunks by the recording software. Since there were no specific light on/off times, the first and last chunks for analysis were chosen based on containing at least 10 minutes of continuous sleep in the second half for the first chunk and the first half for the last chunk. REM periods following sleep staging according to standardized criteria (Berry et al., 2012) were selected for further analyses. REM periods with (phasic REM) and without (tonic REM) eye movements were identified by visual inspection using the EOG channels. Four-second-long segments were coded as phasic when the bipolar EOG channel exhibited at least two consecutive eye movements and featured 100 µV (or larger) amplitudes within the specific time window. Four-second-long segments without significant bursts of eye movements (EOG deflections lower than 25 µV) were categorized as tonic REM. The selection of segments were conducted by research assistants trained in sleep scoring, and the selected 4-s-long periods were visually inspected by a trained sleep researcher to exclude segments with inaccurate categorizations. Segments with artifacts in the ANT and/or ECG channel and those showing inter-ictal spikes or signs of pathological activity were discarded after further visual inspection. To compare REM microstates with resting wakefulness, 5-10 minutes of segments containing eyes-closed wakefulness were selected for further analyses. Since resting state recordings were limited in number in our sample, we selected awake segments from both presleep wakefulness and from periods of wake after sleep onset throughout the night. Preprocessing steps of wake trials were identical to those performed on REM trials. Segments were band pass-filtered between 0.3 and 70 Hz (with Butterworth, zero phase forward and reverse digital filters) and downsampled to 512 Hz.

R-peaks of the ECG channel were identified semi-automatically using scripts of the HEPLAB toolbox (Perakakis, 2019). Available ANT channels in each participant were segmented to epochs of 1000 ms spanning between -200 ms and +800 ms time-locked to the R peak of the ECG channel, yielding to an average number of 302.9 trials (range: 67–744) in phasic, 655.5 trials (range: 83–1263) in tonic, and 378.7 trials (range: 148–614) in resting wakefulness. Sleep architecture and the amount of selected trials for each participant are summarized in **Table2**.

**Table 2.** Sleep architecture of the night records of the subjects involved in the study (all measures are expressed in minutes)

| Patient | Total recording time | TST | WASO | Wake | Stage1 | Stage2 | Stage3 | REM | Phasic REM # | Tonic REM # | Resting wake # |
| --- | --- | --- | --- | --- | --- | --- | --- | --- | --- | --- | --- |
| #1 | 690.3 | 416 | 187 | 274.6 | 59.3 | 226.7 | 38.7 | 91.3 | 252 | 191 | 407 |
| #2 | 596 | 542.3 | 29.6 | 53.7 | 45.3 | 312 | 66.7 | 118.3 | 118 | 337 | 205 |
| #3 | 662.6 | 247.7 | 92.3 | 415 | 62 | 146.3 | 0 | 39.3 | 67 | 83 | 362 |
| #4 | 503 | 461.3 | 38.7 | 41.7 | 10 | 296.7 | 86 | 68.7 | 390 | 1263 | 394 |
| #5 | 450 | 397 | 29.3 | 53 | 24 | 168 | 82.7 | 122.3 | 455 | 708 | 374 |
| #6 | 450 | 347.3 | 76.3 | 102.7 | 26.3 | 274.3 | 14.3 | 32.3 | 113 | 189 | 148 |
| #7 | 450 | 294.3 | 155.6 | 155.7 | 34 | 174.3 | 20 | 66 | 480 | 584 | 407 |
| #8 | 630 | 522 | 88.7 | 108 | 19 | 420.7 | 16.7 | 65.7 | 230 | 1161 | 401 |
| #9 | 720 | 574.3 | 56.7 | 145.7 | 15.7 | 433.3 | 53.3 | 72 | 744 | 1097 | 614 |
| #10 | 630 | 493.3 | 119.3 | 136.7 | 25.7 | 386.3 | 36.3 | 45 | 319 | 782 | 475 |
| #11 | 600 | 560 | 30 | 40 | 47 | 295.7 | 98 | 119.3 | 164 | 819 | 379 |

Phasic and Tonic REM and Awake trial numbers refer to the amount of analyzed 1 sec long segments locked (between -200 and 800 ms) with reference to the R peak of the ECG signal. WASO: Wake after sleep onset refers to wake time after the first epoch of not S1 sleep. TST: Total Sleep Time. Since there were no explicit schedules of lights off, we did not analyze sleep latency and sleep efficiency.

### Data analyses

All analyses were performed in MATLAB, using custom-based scripts and functions from the Fieldtrip toolbox (Oostenveld et al., 2011).

*Heartbeat-evoked potentials* (HEPs) were computed from the ANT recordings. One-second trials, locked to the cardiac signal (from -200 to +800 ms relative to the R-peak), were averaged for each condition (phasic REM, tonic REM, wake) in each participant. For participants with multiple ANT contacts (N = 5), polarity reversals were visually inspected and inverted before averaging over channels within participants. Averaged amplitudes locked to the heartbeat were baseline corrected with their mean amplitudes between -200 and -50 ms to avoid the influence of the rising edge of the R wave (Coll et al., 2021; Schandry et al., 1986). Amplitude fluctuations locked to the heartbeat were statistically compared between conditions along the time window between 350 and 650 ms after the R peak. We focused on this time window because HEP-like waveforms were previously identified in REM sleep at late components (Perogamvros et al., 2019), the influence of the cardiac R and T waves might be reduced at 350 ms after the R-peak (Park and Blanke, 2019), and a previous scalp-EEG study identified HEP differences between REM microstates and wakefulness also in late components (Simor et al., 2021a). To ensure that the observed differences in HEP amplitudes were not caused by differences in ECG activity, the ECG waveforms pertaining to the selected trials were also averaged and compared across the three conditions. The aim of this approach was to control for any confounding influence of ECG activity on the observed differences in HEP across the three conditions (Park and Blanke, 2019).

We performed *time-frequency analyses (TFA)* to characterize heartbeat-related anterothalamic local field potentials in more detail. Power values along the time and frequency dimensions were extracted using Fast Fourier Transformation (FFT) of overlapping, Hanning-tapered, 150-ms long time windows, sliding in 10-ms steps over the selected 1-second long trials. Zero-padding was applied to extend the signal to 1.2 seconds before performing the FFT. Power values between 7 and 45 Hz (with 1 Hz frequency resolution) locked to the R-peak were considered for further analyses. We applied baseline correction (using the period between -200 ms – -50 ms with reference to the R-peak), and changes in power were defined relative to this baseline. The trials of time-frequency power modulations locked to the R-peak were averaged within each condition in each participant. Similarly, to the HEP analysis, the power values of participants with multiple ANT channels were averaged before statistical comparisons.

Finally, *inter-trial phase coherence (ITPC)* was computed to investigate the consistency of the phase of frequency-specific oscillatory activity relative to the heartbeat. ITPC is a measure of the phase consistency of oscillatory signals across trials and is particularly useful for examining whether oscillatory activity is phase-locked to an event, in this case, the R-peak of the heartbeat. The phase information was extracted from the same time-frequency decomposition used for the time-frequency analysis (TFA), which involved applying a Fast Fourier Transformation (FFT) to overlapping, Hanning-tapered, 150-ms time windows sliding in 10-ms steps across the 1-second trials. For each trial, frequency-specific phase angles were computed across the frequencies of interest (7 to 45 Hz with 1 Hz resolution). The padding was set to 1.2 seconds to avoid edge effects and to ensure accurate phase estimation. ITPC was calculated as follows:

The phase and amplitude of the signal at each frequency and time point were extracted using complex values obtained from the FFT. The phase angle (*θ*_*f*,_) at each frequency (*f*) and time point (*t*) was used for subsequent ITPC calculations. ITPC was computed as the absolute value of the average of the unit vectors representing the phase angles across trials:

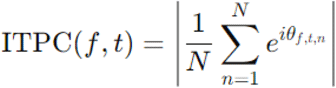

Where *N* is the number of trials, and *θ*_*f, t, n*_ is the phase at *n*th trial at frequency *f* and time *t*. This results in ITPC values ranging from 0 to 1, where 0 indicates completely random phases across trials (i.e., high variability) and 1 indicates perfect phase alignment (i.e., high consistency)(Cohen, 2014).

ITPC values were baseline corrected (baseline between: -200 ms and -50 ms) to highlight relative changes in phase consistency with respect to heartbeats. For TFA and ITPC, statistical comparisons between conditions (phasic REM, tonic REM, wakefulness) were performed within the time range of 50 ms to 650 ms after the R-peak, as amplitude and phase modulations in intracranial recordings were observed over a broader time range including also early components (García-Cordero et al., 2017; Park et al., 2018).

### Statistical analyses

Differences between conditions (phasic REM, tonic REM, wakefulness) were evaluated by pairwise comparisons using cluster-based nonparametric statistics, a suitable approach for the statistical analyses of EEG data. Cluster-based permutation statistics does not require assumptions of data distribution and it efficiently addresses the issue of multiple comparisons (Maris and Oostenveld, 2007). In brief, two-tailed, paired t-tests were performed contrasting each condition (i.e. phasic vs. tonic, phasic vs. wake, tonic vs. wake) along each data point within the time axis (in the case of HEP), and time x frequency axes (in the case of TFA and ITPC). Clusters were defined if at least two adjacent time points or neighbouring frequency bins showed significant differences at the alpha level below .05. These clusters were selected to compute the observed cluster statistic defined by the sum of all the *t*-values that formed a given cluster. The same process was repeated 1000 times by randomly shuffling conditions (using MonteCarlo simulations). From these simulations the largest clusters were extracted in order to create a distribution of the maximal clusters produced by chance. Finally, the observed cluster statistics were tested (with an alpha value of .05) against the probability distribution of the largest simulated clusters.

## Results

### Anterior thalamic HEPs are different in phasic REM compared to tonic REM and wakefulness

First, we examined whether HEPs (heart-evoked potentials) differed across the three states within the time range of interest (350–650 ms) (**Fig1A**). Significant positive clusters emerged when comparing phasic REM to tonic REM (424.3–529.9 ms, t_maxsum_ = 145.9478, cluster-level p = 0.0151; 571–600.4 ms, t_maxsum_ = 38.5384, cluster-level p = 0.0493) and when comparing phasic REM to wakefulness (373.4–488.8 ms, t_maxsum_ = 144.2944, cluster-level p = 0.122; 500.6–514.3 ms, t_maxsum_ = 19.5978, cluster-level p = 0.474; 541.7–555.4 ms, t_maxsum_ = 18.2176, cluster-level p = 0.0474). However, no significant clusters were observed when contrasting HEPs between tonic REM and wakefulness.

**Fig 1.**
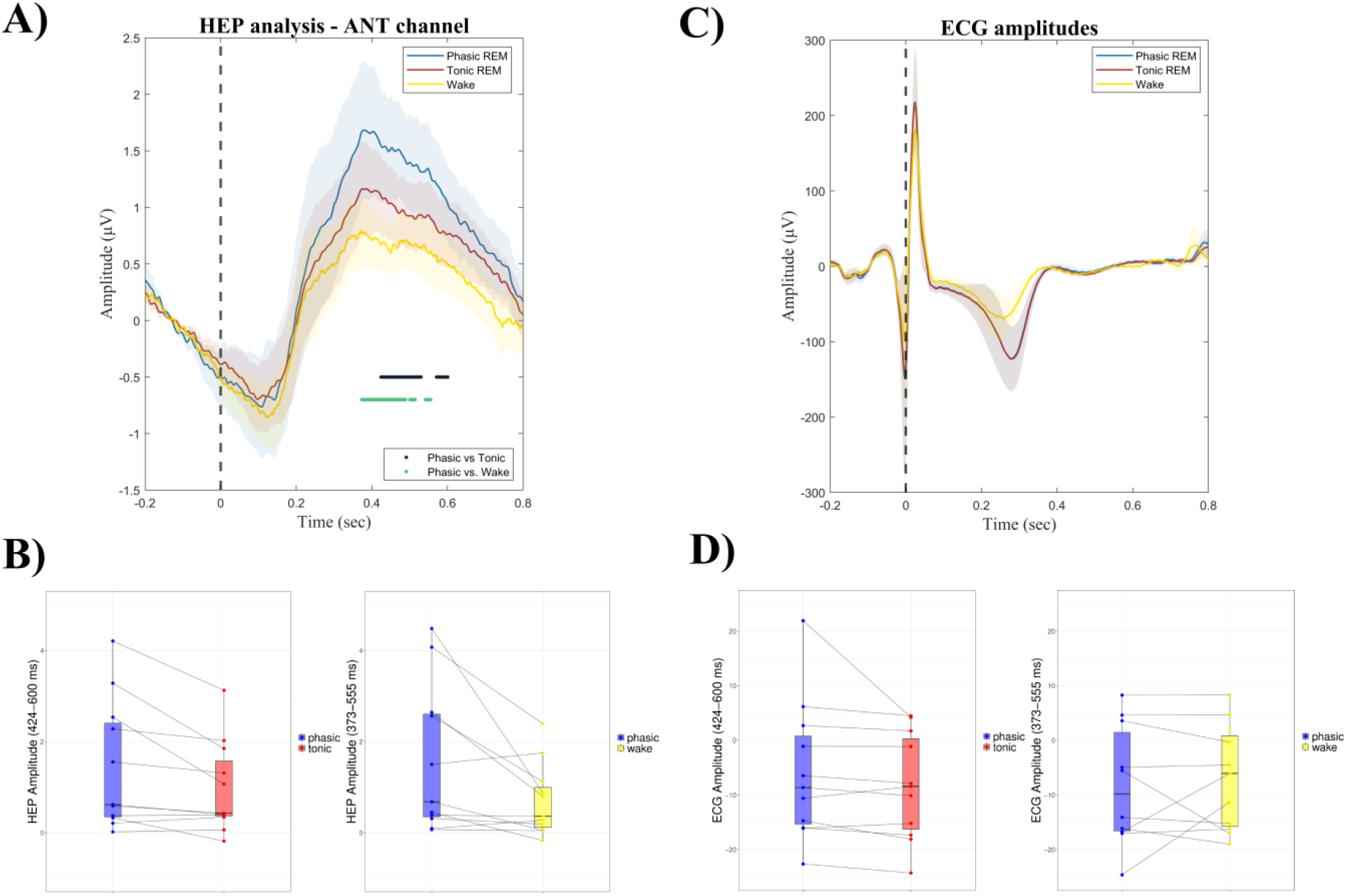
Heartbeat evoked potential (HEP) in the anterior thalamus in phasic and tonic REM sleep and resting wakefulness. A) HEP amplitudes in phasic and tonic REM and wakefulness. HEPs differed at late potentials (over ∼400 and 600 ms) between phasic REM and tonic REM, as well as between phasic REM and wakefulness. Black and green vertical lines indicate significant differences along the time axis between phasic and tonic REM, and phasic REM and wakefulness, respectively. The vertical dashed line mark the time point of the R-peak. B) HEP amplitudes averaged along the time ranges where significant differences emerged across conditions. C) Averaged ECG amplitudes in the three conditions. No significant differences were observed between the three vigilance states in ECG amplitudes. D) ECG amplitudes averaged along the time ranges where significant HEP differences emerged between conditions. No significant differences in ECG amplitudes over the time range of interest emerged, indicating that HEP differences were not confounded by ECG activity.

To control for potential confounding effects of cardiac activity, we conducted pairwise comparisons of ECG amplitudes between phasic and tonic REM, as well as between phasic REM and wakefulness, within the same time range (350–650 ms) (**Fig1B**). We also examined whether mean ECG amplitudes differed in the specific time ranges where significant HEP clusters were observed. The ECG amplitudes did not significantly differ between phasic and tonic REM (t(10) = 1.518, p = 0.160, Cohen’s d = 0.458) or between phasic REM and wakefulness (t(10) = 0.606, p = 0.558, Cohen’s d = 0.183) during these periods (see **Fig1CD**). Additionally, no significant ECG amplitude clusters were found across conditions between 350 and 650 ms.

These analyses suggest that HEPs in the ANT during phasic REM periods differ significantly from those during tonic REM and resting wakefulness (see **Fig1**).

### Reduced heartbeat-related anterior thalamic fast frequency power in phasic REM

Next, we explored power changes relative to the R-peak, and examined time x frequency power modulations across the three conditions. Similarly to HEPs, significant clusters were observed between phasic and tonic REM (time range: 315.1–444.2 ms, frequency range: 18.3–35 Hz, t_maxsum_ = -409.3936, cluster level p = 0.036), and between phasic REM and wakefulness (time range: 164.4–465.8 ms, frequency range: 19.2–45 Hz, t_maxsum_ = -1276.6199, cluster level p = 0.002 and time range: 524.5–634.1 ms, frequency range: 28.3–45 Hz, t_maxsum_ = -322.3900, cluster level p = 0.048), while the comparison between tonic REM and wakefulness did not yield significant clusters. As shown in **Fig2**, fast frequency activity in the beta and low gamma range (∼20–40 Hz) locked to the R-peak was relatively reduced in phasic REM as compared to wakefulness and tonic REM sleep. Although fast frequency power modulation locked to the R-peak seemed to peak at relatively higher frequencies in wakefulness (above 30 Hz) than in tonic REM (between 20 and 30 Hz), these differences were not significant after cluster-based correction.

**Fig 2.**
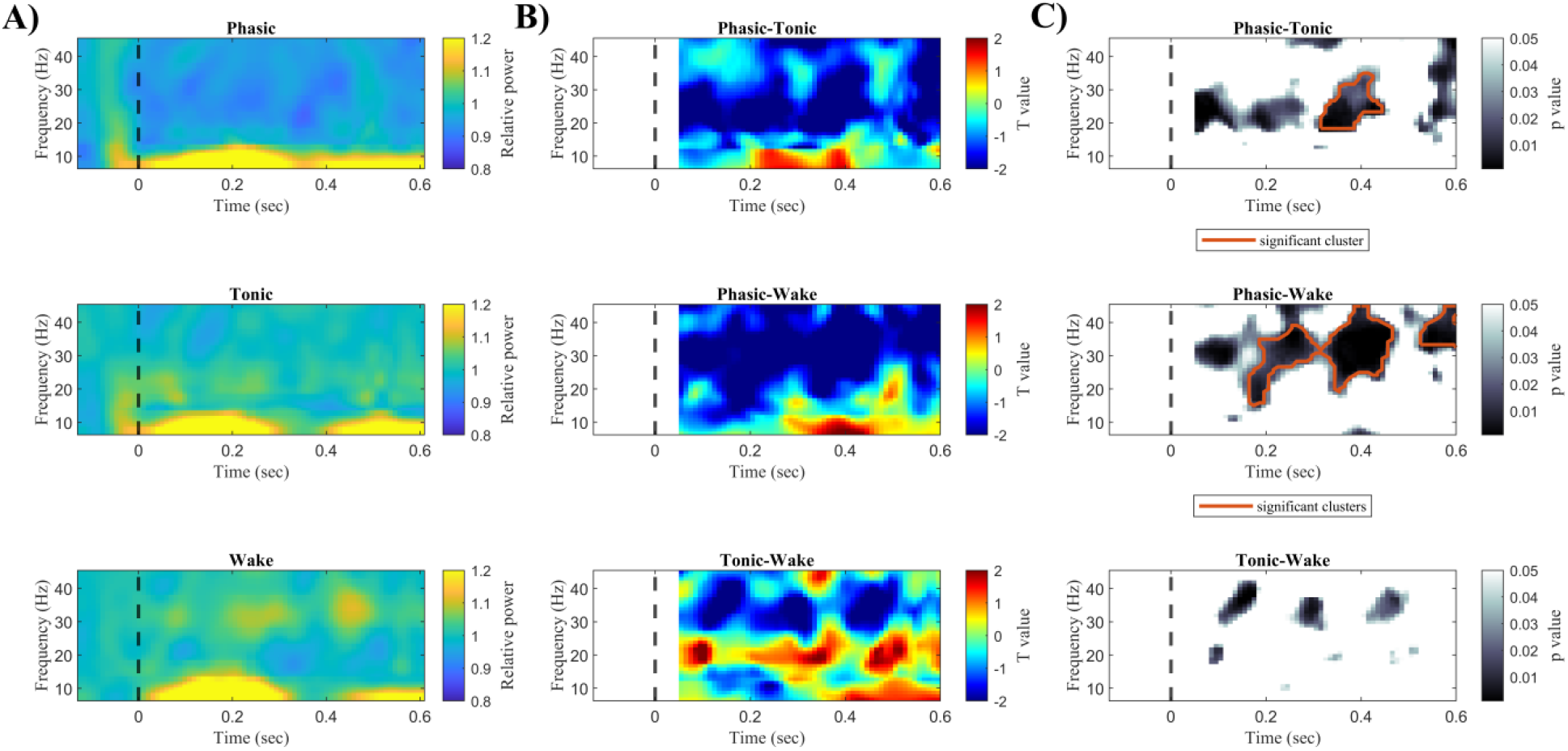
Power modulations in the anterior thalamus locked to the R-peak of the ECG signal. A) Changes in time frequency power relative to the baseline period (between -200 and -50ms) are visualized for phasic REM, tonic REM, and wakefulness. Values above 1 indicate increased power, whereas values below 1 mean reductions in power relative to the baseline. B) Pairwise statistical comparisons across conditions highlighting the value of the statistical test (T value) along the time and frequency axis. C) Uncorrected significant differences (p<0.05) at each time x frequency point and significant clusters remaining after the correction for multiple comparisons are visualized (with red contour). Power changes locked to the R-peak in the beta and low gamma range (∼20–40 Hz) were relatively reduced in phasic REM as compared to wakefulness and tonic REM sleep. The vertical dashed line mark the time point of the R-peak.

To explore, if differences across conditions in heartbeat-related high frequency activity were mainly driven by a relative reduction of beta-low gamma power in phasic REM, or a relative increase in high frequency power in tonic REM and wake, we contrasted power values of the baseline periods with those of the periods after the R-peak within each condition. To test whether power in the beta and low gamma (20–40 Hz) frequency range differed between the baseline (−200 ms to -50 ms) and the post-R-peak periods, we contrasted the 150 ms-long baseline (pre-R-peak) periods with the subsequent period after the R-peak chunked into four non-overlapping 150ms-long segments between 50 and 650 ms. Fast (20–40 Hz) frequency power in the baseline periods were hence compared with the post-R-peak period using cluster-based permutations statistics along the time axis. As shown in **Fig3**, high frequency power fluctuations as referenced to the baseline (pre-R-peak period) were relatively reduced in phasic REM, but not in tonic REM and wakefulness. Statistically significant differences between the baseline and post-R-peak periods were only observed in phasic REM (STAT parameters: time range: 324.8–354.2 ms, t_maxsum_ = -9.7095, cluster level p = 0.0068, and time range: 375.7–434.4 ms, t_maxsum_ = -20.9739, cluster level p = 0.0020 and time range: 565.5–575.3 ms, t_maxsum_ = -5.3738, cluster level p = 0.0283), but not in tonic REM and wakefulness.

**Fig 3.**
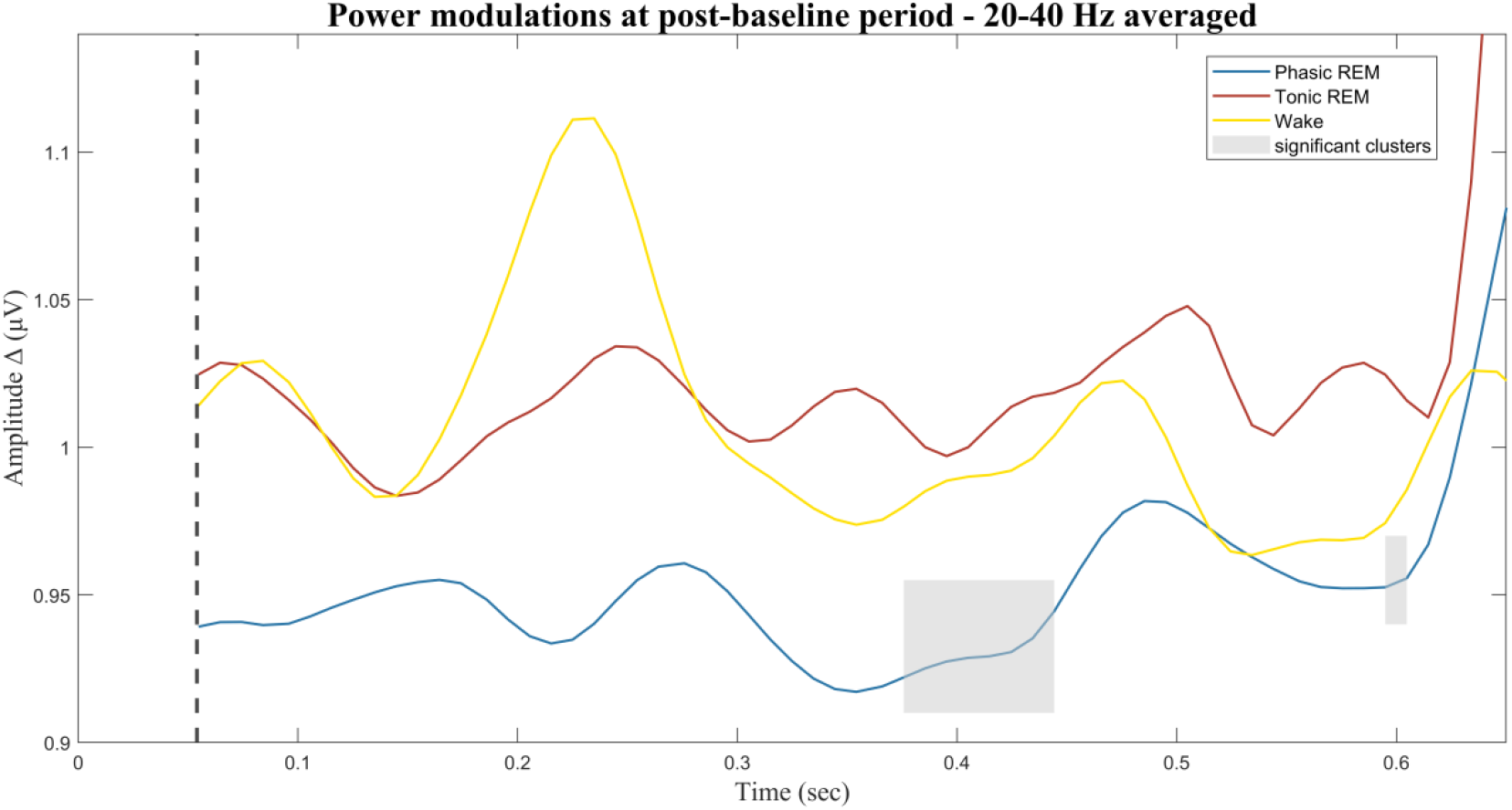
Fast-frequency power changes in the anterior thalamus locked to the R-peak. Power changes relative to the baseline period (−200 and -50 ms) between 20-40 Hz were averaged and visualized between 50 ms and 650 ms. Values above 1 indicate increased power, whereas values below 1 mean reductions in power relative to the baseline. High frequency activity was significantly relatively reduced in phasic REM after the R-peak at time ranges marked by the grey shaded bars. Post-baseline periods did not significantly differ from baseline periods in tonic REM and wakefulness. The dashed vertical line mark the start of the analyzed post-baseline period (+50 ms).

### Heartbeat-related low-frequency phase modulation in the anterior thalamus across all vigilance states

Pairwise comparisons of ITPC between REM microstates and wakefulness are presented in **Fig4**. As indicated in the figure, no significant differences emerged between conditions after cluster-based correction of multiple comparisons. On an exploratory level, uncorrected t-tests showed trend-like differences between phasic and tonic REM microstates, showing increased inter-trial phase coherence in lower (∼ 10–17 Hz) and gamma (∼40–45 Hz) frequencies during tonic REM, and relatively increased phase coherence in the beta (∼ 20–30 Hz) frequency range during phasic REM. Nominal differences between phasic REM and wakefulness showed a similar pattern with respect to the relative increase in 20–30 Hz ITPC in phasic REM, and the relative increase in 40–45 Hz gamma ITPC in wakefulness. ITPC in tonic REM showed slightly increased values in the (8–15 Hz) alpha frequency range compared to wakefulness. Nevertheless, none of the above differences remained significant after correction for multiple comparisons.

**Fig 4.**
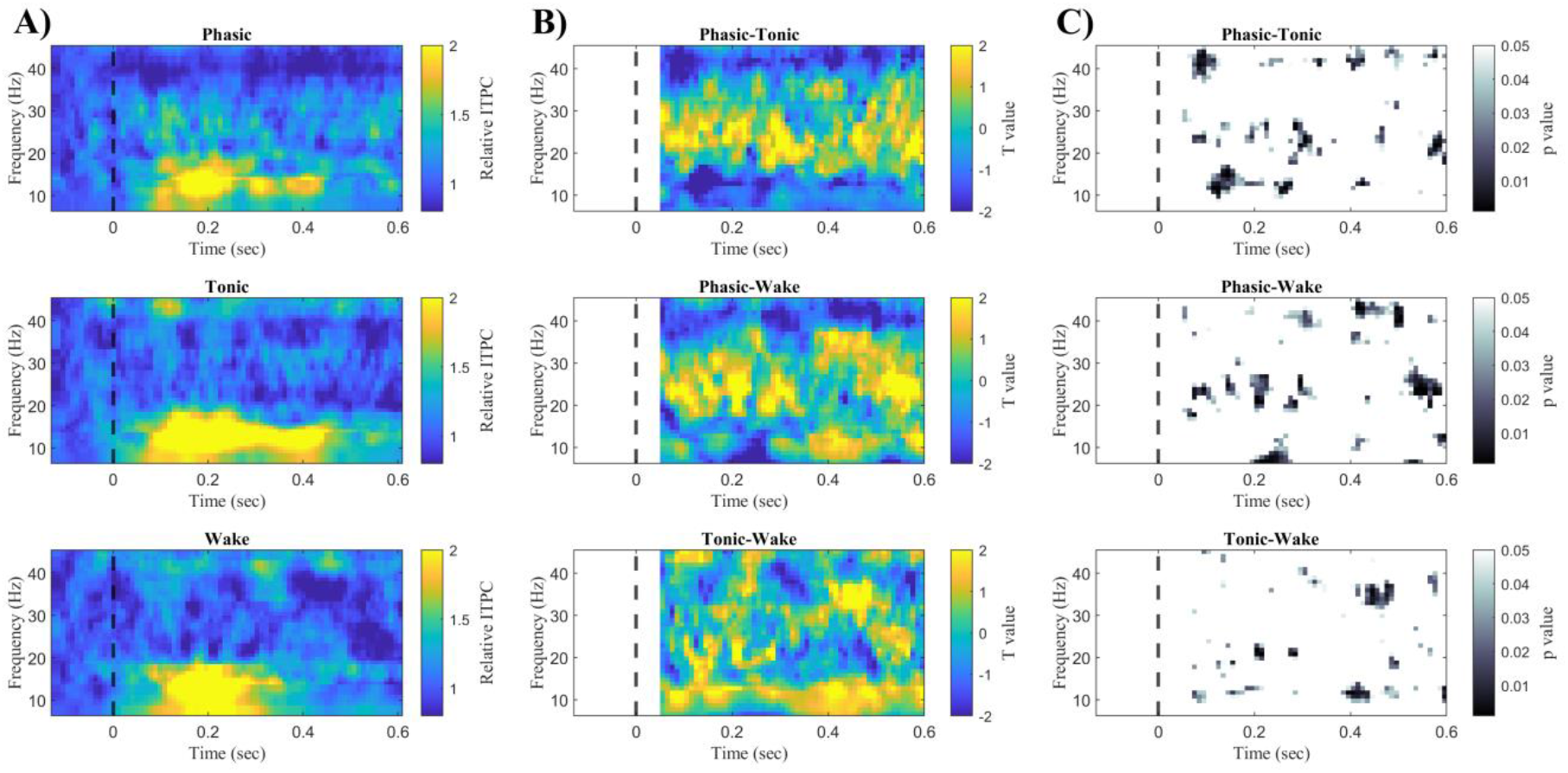
Inter-trial phase coherence (ITPC) in the anterior thalamus relative to the R-peak of the ECG signal. A) Changes in ITPC relative to the baseline period (between -200 and -50ms) are visualized for phasic REM, tonic REM, and wakefulness. Values above 1 indicate increased power, whereas values below 1 mean reductions in power relative to the baseline. B) Pairwise statistical comparisons of ITPC changes across conditions highlighting the value of the statistical test (T value) along the time and frequency axis. C) Uncorrected significant differences (p<0.05) at each time x frequency point are visualized. Changes in ITPC locked to the R-peak were not significantly different across conditions.

Given that ITPC analyses indicated a robust heartbeat-related increase in low-frequency phase coherence between 7–20 Hz in all of the conditions (see **Fig4**), we examined if such increase was statistically significant as compared to ITPC values of the baseline (pre-R-peak) period. Therefore, ITPC values at baseline (from - 200 ms to -50 ms) were statistically compared (applying cluster-based permutation statistics) with post-R periods chunked into four non-overlapping 150ms-long segments between 50 and 650 ms. ITPC between ∼ 7–20 Hz showed robust increases in all conditions after the R-peak between 130–444 ms in phasic REM (t_maxsum range_: 182-233, cluster level p_range:_ 0.0001-0.01), between 54–444 ms in tonic REM (t_maxsum range_: 139– 258, cluster level p_range:_ 0.001-0.03), and between 55–424 ms in wakefulness (**Fig5**). In addition, a negative cluster emerged in phasic REM, pointing to decreased ITPC at 36–45 Hz between 475–524ms after the R-peak (t_maxsum_ = -81.1285, cluster level p = 0.0107).

**Fig 5.**
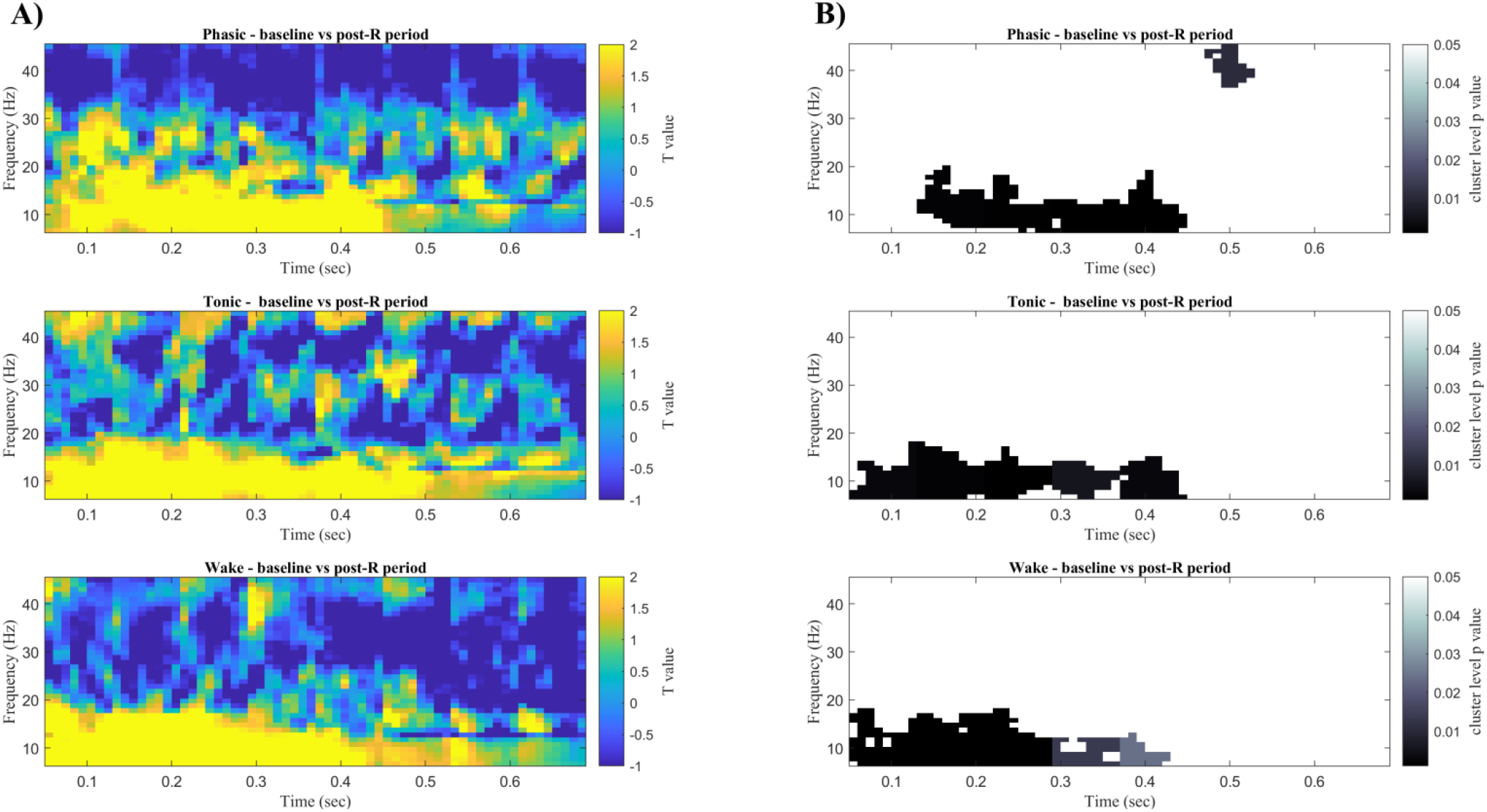
Changes in inter-trial coherence (ITPC) in the anterior thalamus locked to the R-peak. A) Statistical comparisons of ITPC between the baseline period (−200 – -50 ms) and the post-baseline periods (50 – 650 ms) in phasic REM, tonic REM, and wakefulness. Statistical parameters (T value) of within-state comparisons along the time x frequency axis are highlighted. B) Significant clusters at each time x frequency point after cluster-based permutation statistics are highlighted.

## Discussion

We studied interoceptive processing during REM sleep and resting wakefulness by measuring local field potentials in the anterior nucleus of the thalamus (ANT) locked to the heartbeat. Heartbeat-related neural activity in the ANT was distinct during REM periods with ocular activity (phasic REM) compared to tonic REM and wakefulness. More specifically, evoked potentials and power fluctuations of beta–low gamma frequencies locked to the heartbeat were significantly different in phasic REM states compared to tonic REM and wakefulness. On the other hand, heartbeat-related amplitude and power fluctuations did not show significant differences between tonic REM and wakefulness. Moreover, heartbeat-related neural signals exhibited pronounced phase synchronization at lower (7–20 Hz) oscillatory activity in the ANT lasting 400 ms after the R-peak in all vigilance states, but reduced gamma synchronization at later time points (∼500 ms after the R-peak) in phasic REM only. Our findings indicate that heartbeat-related neural activity is detectable at the level of the ANT, showing distinct signatures of interoceptive processing in phasic REM compared to tonic REM and wakefulness.

Sleep in general, and especially REM sleep is usually viewed as a state of sensory disconnection; however, a wide variety of studies indicate that the sleeping brain is not fully impermeable to external stimulation (Salvesen et al., 2024). Whereas cortical reactivity to external inputs in the sleeping brain were extensively studied (Bastuji and García-Larrea, 1999; Colrain, 2005; Halasz et al., 2004; Halász et al., 2014), neural responses to interoceptive signals received relatively less attention. Nonetheless, previous studies indicate that beyond exteroception (mostly studied in the auditory domain), the sleeping brain is also susceptible to interoceptive signals, which, by providing a rich source of information, seem to affect the regulation of sleep and arousal (Mazza et al., 2012; Wei and Van Someren, 2020).

Recent studies suggest that during REM sleep, the alternation between phasic and tonic periods facilitates environmental disconnection and alertness, respectively. Specifically, while the processing of auditory inputs is largely reduced in phasic REM sleep, cortical responses to external stimulation are reinstated during tonic REM periods (Koroma et al., 2020; Sallinen et al., 1996; Takahara et al., 2006, 2002; Wehrle et al., 2007). In line with studies on auditory processing, a recent study (Simor et al., 2021a) examined heartbeat-evoked potentials (HEPs) of scalp electrode recordings in phasic and tonic REM sleep, and observed different modulation of HEPs at late components (∼550–650 ms) in phasic compared to tonic REM and wakefulness. Moreover, similar to results in auditory stimulation studies (Sallinen et al., 1996; Wehrle et al., 2007), HEPs in tonic REM sleep featured intermediate values between those of phasic REM and wakefulness (Simor et al., 2021a). Our findings in the ANT align with this pattern: amplitude modulations locked to the heartbeat in the ANT differed between phasic and tonic REM during late potentials (400–600 ms) and between phasic REM and wakefulness (370–550 ms), with HEPs in tonic REM appearing intermediate between phasic REM and wakefulness. Moreover, such differences in HEPs were not confounded by differences in ECG amplitudes. Notably, HEP modulations differing across states were observed at slightly earlier time points in the current study compared to findings on the level of the scalp. Since the ANT is a critical site in the transmission of subcortical signals to cortical regions, we may speculate that signals from the visceral pathway reach the ANT before being processed in cortical layers. Likewise, heartbeat-related neural responses in deeper brain regions (e.g. insula) were observed at relatively earlier time points (Park et al., 2018) than HEPs in scalp EEG recordings (Coll et al., 2021).

Heartbeat-related phase modulations of oscillatory activity, as quantified by ITPC, revealed a robust increase in phase alignment at lower frequencies (7–20 Hz) during REM states and wakefulness. This finding aligns with a previous study that examined ITPC of LFPs recorded intracranially from the insula (Park et al., 2018), a key region involved in interoceptive processing (Berntson and Khalsa, 2021; Chen et al., 2021; Craig and Craig, 2009; Park and Blanke, 2019; Seth, 2013). In that study, the authors observed an increase in ITPC lasting approximately 400 ms after the R-peak during resting wakefulness and during conditions that enhance interoceptive processing (Park et al., 2018). Our findings corroborate this result by demonstrating strikingly similar increases in ITPC with respect to both temporal and frequency characteristics. Moreover, our finding indicates that oscillatory activity in the ANT seems to track cardiac signals not only in wakefulness, but also in REM sleep. While phase alignment in lower frequencies were not distinct between the three vigilance states, beta–low gamma power was relatively reduced in phasic REM compared to tonic REM and resting wakefulness. The reduction of heartbeat-related beta–low gamma power was paralleled by a decrease in gamma synchronization (ITPC) apparent only in phasic REM sleep. In sum, in addition to HEPs, modulations of time-frequency power and synchronization in the beta–low gamma and gamma frequency range, respectively, point to distinct processing of cardiac signals in the ANT during the phasic REM state.

Although the specific role of heartbeat-related activity in the ANT in interoception should be explored in future studies that experimentally manipulate interoceptive processing, our findings offer new insights into the distinctive nature of phasic REM sleep. We propose that reduced high frequency power and synchronization in response to heartbeats may reflect the attenuation of interoceptive afferents during phasic REM periods. Beta EEG oscillations are commonly implicated in sensorimotor processing (Pfurtscheller and Da Silva, 1999). Recent evidence suggests that beta EEG waves are inherently transient in nature, whereas distinct types of beta bursts have different functions. Indeed, according to current insights into the functional role of this oscillatory bursts, decreased beta activity in the period after the ECG R-peak in phasic REM sleep could indicate blunted interareal communication (Lundqvist et al., 2024), which might arise shortly after the interoceptive input. In addition, gamma band activity is associated with cortical information processing, modulating attention and awareness (Fell et al., 2003; Kaiser and Lutzenberger, 2003), and enhancing the neural gain of exteroceptive (van Es and Schoffelen, 2019) and interoceptive inputs (Li et al., 2023). For instance, induced changes (increases) in gamma band power were linked to selective attention to nociceptive stimuli (Tiemann et al., 2010) and to the subjective intensity of pain (Zhang et al., 2012). Phasic REM sleep is viewed as a closed-loop state in which the influence of (bottom-up) lower-level signals arising from the sensory periphery is largely reduced (Simor et al., 2020; Wehrle et al., 2007). The attenuation of cardiac signals in phasic REM sleep might be associated with reduced neural gain on lower-level afferent pathways. This diminished neural gain on interoceptive signals is consistent with the particularly unstable nature of autonomic nervous system activity during phasic REM sleep, characterized by irregular breathing, sudden surges in heart rate and blood pressure, and impaired thermoregulation (Amici and Zoccoli, 2021). Accordingly, even autonomic reflexes that facilitate homeostatic adjustments, such as the baroreflex, have been observed to be suppressed by central commands during the phasic period of REM sleep (Silvani, 2008; Silvani and Dampney, 2013). Disconnection from the sensory periphery during periods of phasic REM sleep is paralleled by increased activity in a thalamocortical network which seems to operate in isolation from the external environment (Maquet et al., 1996; Miyauchi et al., 2009; Wehrle et al., 2007). Although the functions of such internally driven processing remain to be elusive, recent animal studies suggest that saccades of REMs are associated with the internal representation of a virtual environment (i.e. the generative model of the real world) in which the animal simulate actions and sensory outcomes (Senzai and Scanziani, 2024, 2022), which may give rise to perceptually immersive dreams.

Acknowledging the limitations of the present study, we note that due to the small sample size, our findings should be interpreted with caution. Further research is needed to determine whether distinct heartbeat-related responses in phasic REM are present in other cortical regions involved in interoception (e.g., the insula and somatosensory cortex). Additionally, sleep recordings were conducted after surgery, leading to fragmented sleep and reduced REM sleep in our sample. Therefore, we cannot fully rule out the possibility that the medical environment affected sleep quality and interoceptive processes. Finally, the patients were on antiseizure medications, which may have influenced oscillatory activity.

Despite these limitations, our sample offered a unique opportunity to examine heartbeat-related activity in the anterior thalamus. Our findings provided novel empirical evidence to the understanding of REM sleep heterogeneity and highlighted the distinct cognitive processing that occurs in phasic REM periods, particularly in the context of interoception.

## Acknowledgements

The work was supported by the National Research, Development and Innovation Office of Hungary K_128117 (RB), FK 142945 (PS) and by the Ministry of Innovation and Technology of Hungary (TKP2021-EGA-25, TKP2021-NKTA-47). PS received the Janos Bolyai scholarship of the Hungarian Academy of Sciences. ZR was Supported by the ÚNKP-23-1 New National Excellence Program of the Ministry for Innovation and Technology from the source of the National Research, Development and Innovation Fund.

## References

Amici, R., Zoccoli, G., 2021. Physiological Changes in the Autonomic Nervous System During Sleep, in: Chokroverty, S., Cortelli, P. (Eds.), Autonomic Nervous System and Sleep. Springer International Publishing, Cham, pp. 43–50. 10.1007/978-3-030-62263-3_5

Bastuji, H., García-Larrea, L., 1999. Evoked potentials as a tool for the investigation of human sleep. Sleep medicine reviews 3, 23–45.

Bastuji, H., Perchet, C., Legrain, V., Montes, C., Garcia-Larrea, L., 2008. Laser evoked responses to painful stimulation persist during sleep and predict subsequent arousals. PAIN(r) 137, 589–599.

Berntson, G.G., Khalsa, S.S., 2021. Neural Circuits of Interoception. Trends in Neurosciences 44, 17–28. 10.1016/j.tins.2020.09.011

Berry, R.B., Budhiraja, R., Gottlieb, D.J., Gozal, D., Iber, C., Kapur, V.K., Marcus, C.L., Mehra, R., Parthasarathy, S., Quan, S.F., 2012. Rules for scoring respiratory events in sleep: update of the 2007 AASM manual for the scoring of sleep and associated events: deliberations of the sleep apnea definitions task force of the American Academy of Sleep Medicine. Journal of clinical sleep medicine 8, 597–619.

Blumberg, M.S., Coleman, C.M., Gerth, A.I., McMurray, B., 2013. Spatiotemporal Structure of REM Sleep Twitching Reveals Developmental Origins of Motor Synergies. Current Biology 23, 2100–2109. 10.1016/j.cub.2013.08.055

Blumberg, M.S., Lesku, J.A., Libourel, P.-A., Schmidt, M.H., Rattenborg, N.C., 2020. What Is REM Sleep? Current Biology 30, R38–R49. 10.1016/j.cub.2019.11.045

Boccia, M., Teghil, A., Raimo, S., Di Vita, A., Grossi, D., Guariglia, C., Palermo, L., 2023. Neural substrates of interoceptive sensibility: An integrated study in normal and pathological functioning. Neuropsychologia 183, 108504. 10.1016/j.neuropsychologia.2023.108504

Bueno-Junior, L.S., Ruckstuhl, M.S., Lim, M.M., Watson, B.O., 2023. The temporal structure of REM sleep shows minute-scale fluctuations across brain and body in mice and humans. Proc. Natl. Acad. Sci. U.S.A. 120, e2213438120. 10.1073/pnas.2213438120

Carskadon, M.A., Dement, W.C., 2005. Normal human sleep: an overview. Principles and practice of sleep medicine 4, 13–23.

Chen, W.G., Schloesser, D., Arensdorf, A.M., Simmons, J.M., Cui, C., Valentino, R., Gnadt, J.W., Nielsen, L., Hillaire-Clarke, C.S., Spruance, V., 2021. The emerging science of interoception: sensing, integrating, interpreting, and regulating signals within the self. Trends in neurosciences 44, 3–16.

Child, N.D., Benarroch, E.E., 2013. Anterior nucleus of the thalamus: Functional organization and clinical implications. Neurology 81, 1869–1876. 10.1212/01.wnl.0000436078.95856.56

Cohen, M.X., 2014. Analyzing Neural Time Series Data: Theory and Practice. The MIT Press. 10.7551/mitpress/9609.001.0001

Coll, M.-P., Hobson, H., Bird, G., Murphy, J., 2021. Systematic review and meta-analysis of the relationship between the heartbeat-evoked potential and interoception. Neuroscience & Biobehavioral Reviews 122, 190–200.

Colrain, I.M., 2005. The K-Complex: A 7-Decade History. Sleep 28, 255–273. 10.1093/sleep/28.2.255

Craig, Arthur D., Craig, A. D., 2009. How do you feel–now? The anterior insula and human awareness. Nature reviews neuroscience 10.

Datta, S., 2004. Activation of Phasic Pontine-Wave Generator Prevents Rapid Eye Movement Sleep Deprivation-Induced Learning Impairment in the Rat: A Mechanism for Sleep-Dependent Plasticity. Journal of Neuroscience 24, 1416–1427. 10.1523/JNEUROSCI.4111-03.2004

Ermis, U., Krakow, K., Voss, U., 2010. Arousal thresholds during human tonic and phasic REM sleep. Journal of sleep research 19, 400–406.

Feldman, M.J., Bliss-Moreau, E., Lindquist, K.A., 2024. The neurobiology of interoception and affect. Trends in Cognitive Sciences.

Fell, J., Fernández, G., Klaver, P., Elger, C.E., Fries, P., 2003. Is synchronized neuronal gamma activity relevant for selective attention? Brain Research Reviews 42, 265–272. 10.1016/S0165-0173(03)00178-4

Fernández-Mendoza, J., Lozano, B., Seijo, F., Santamarta-Liébana, E., José Ramos-Platón, M., Vela-Bueno, A., Fernández-González, F., 2009. Evidence of Subthalamic PGO-like Waves During REM Sleep in Humans: a Deep Brain Polysomnographic Study. Sleep 32, 1117–1126. 10.1093/sleep/32.9.1117

García-Cordero, I., Esteves, S., Mikulan, E.P., Hesse, E., Baglivo, F.H., Silva, W., García, M. del C., Vaucheret, E., Ciraolo, C., García, H.S., Adolfi, F., Pietto, M., Herrera, E., Legaz, A., Manes, F., García, A.M., Sigman, M., Bekinschtein, T.A., Ibáñez, A., Sedeño, L., 2017. Attention, in and Out: Scalp-Level and Intracranial EEG Correlates of Interoception and Exteroception. Front. Neurosci. 11, 411. 10.3389/fnins.2017.00411

Gonzalo-Ruiz, A., Sanz-Anquela, M.J., Lieberman, A.R., 1995. Cholinergic projections to the anterior thalamic nuclei in the rat: a combined retrograde tracing and choline acetyl transferase immunohistochemical study. Anat Embryol 192, 335–349. 10.1007/BF00710103

Halász, P., Bódizs, R., Parrino, L., Terzano, M., 2014. Two features of sleep slow waves: homeostatic and reactive aspects–from long term to instant sleep homeostasis. Sleep medicine 15, 1184–1195.

Halasz, P., Terzano, M., Parrino, L., Bodizs, R., 2004. The nature of arousal in sleep. J Sleep Res 13, 1–23. 10.1111/j.1365-2869.2004.00388.x

Hasegawa, E., Miyasaka, A., Sakurai, K., Cherasse, Y., Li, Y., Sakurai, T., 2022. Rapid eye movement sleep is initiated by basolateral amygdala dopamine signaling in mice. Science 375, 994–1000.

Hong, J., Lozano, D.E., Beier, K.T., Chung, S., Weber, F., 2023. Prefrontal cortical regulation of REM sleep. Nature Neuroscience 26, 1820–1832.

Immanuel, S.A., Pamula, Y., Kohler, M., Martin, J., Kennedy, D., Nalivaiko, E., Saint, D.A., Baumert, M., 2014. Heartbeat Evoked Potentials during Sleep and Daytime Behavior in Children with Sleep-disordered Breathing. Am J Respir Crit Care Med 190, 1149–1157. 10.1164/rccm.201405-0920OC

Jouvet, M., 1965. Paradoxical sleep—a study of its nature and mechanisms. Progress in brain research 18, 20–62.

Kaiser, J., Lutzenberger, W., 2003. Induced gamma-band activity and human brain function. Neuroscientist 9, 475–484. 10.1177/1073858403259137

Keil, A., Debener, S., Gratton, G., Junghöfer, M., Kappenman, E.S., Luck, S.J., Luu, P., Miller, G.A., Yee, C.M., 2014. Committee report: publication guidelines and recommendations for studies using electroencephalography and magnetoencephalography. Psychophysiology 51, 1–21.

Koroma, M., Lacaux, C., Andrillon, T., Legendre, G., Léger, D., Kouider, S., 2020. Sleepers selectively suppress informative inputs during rapid eye movements. Current Biology 30, 2411–2417.

Li, W., Ma, L., Yang, G., Gan, W.-B., 2017. REM sleep selectively prunes and maintains new synapses in development and learning. Nature neuroscience 20, 427–437.

Li, Z., Zhang, L., Zeng, Y., Zhao, Q., Hu, L., 2023. Gamma-band oscillations of pain and nociception: A systematic review and meta-analysis of human and rodent studies. Neuroscience & Biobehavioral Reviews 146, 105062. 10.1016/j.neubiorev.2023.105062

Lundqvist, M., Miller, E.K., Nordmark, J., Liljefors, J., Herman, P., 2024. Beta: bursts of cognition. Trends in Cognitive Sciences 28, 662–676. 10.1016/j.tics.2024.03.010

Luppi, P.-H., Clément, O., Sapin, E., Gervasoni, D., Peyron, C., Léger, L., Salvert, D., Fort, P., 2011. The neuronal network responsible for paradoxical sleep and its dysfunctions causing narcolepsy and rapid eye movement (REM) behavior disorder. Sleep medicine reviews 15, 153–163.

Manzano, G.M., Ragazzo, P.C., Tavares, S.M., Marino Jr, R., 1986. Anterior zygomatic electrodes: a special electrode for the study of temporal lobe epilepsy. Stereotactic and Functional Neurosurgery 49, 213–217.

Maquet, P., Péters, J.-M., Aerts, J., Delfiore, G., Degueldre, C., Luxen, A., Franck, G., 1996. Functional neuroanatomy of human rapid-eye-movement sleep and dreaming. Nature 383, 163–166.

Maris, E., Oostenveld, R., 2007. Nonparametric statistical testing of EEG-and MEG-data. Journal of Neuroscience Methods 164, 177–190. 10.1016/j.jneumeth.2007.03.024

Mazza, S., Magnin, M., Bastuji, H., 2012. Pain and sleep: From reaction to action. Neurophysiologie Clinique/Clinical Neurophysiology 42, 337–344. 10.1016/j.neucli.2012.05.003

Miyauchi, S., Misaki, M., Kan, S., Fukunaga, T., Koike, T., 2009. Human brain activity time-locked to rapid eye movements during REM sleep. Exp Brain Res 192, 657–667. 10.1007/s00221-008-1579-2

Oostenveld, R., Fries, P., Maris, E., Schoffelen, J.-M., 2011. FieldTrip: Open Source Software for Advanced Analysis of MEG, EEG, and Invasive Electrophysiological Data. Computational Intelligence and Neuroscience 2011, 1–9. 10.1155/2011/156869

Park, H.-D., Bernasconi, F., Salomon, R., Tallon-Baudry, C., Spinelli, L., Seeck, M., Schaller, K., Blanke, O., 2018. Neural Sources and Underlying Mechanisms of Neural Responses to Heartbeats, and their Role in Bodily Self-consciousness: An Intracranial EEG Study. Cerebral Cortex 28, 2351–2364. 10.1093/cercor/bhx136

Park, H.-D., Blanke, O., 2019. Heartbeat-evoked cortical responses: underlying mechanisms, functional roles, and methodological considerations. Neuroimage 197, 502–511.

Perakakis, P., 2019. HEPLAB: a Matlab graphical interface for the preprocessing of the heartbeat-evoked potential. Zenodo.

Perogamvros, L., Park, H.-D., Bayer, L., Perrault, A.A., Blanke, O., Schwartz, S., 2019. Increased heartbeat-evoked potential during REM sleep in nightmare disorder. NeuroImage: Clinical 22, 101701. 10.1016/j.nicl.2019.101701

Pfurtscheller, G., Da Silva, F.L., 1999. Event-related EEG/MEG synchronization and desynchronization: basic principles. Clinical neurophysiology 110, 1842–1857.

Pollatos, O., Schandry, R., Auer, D.P., Kaufmann, C., 2007. Brain structures mediating cardiovascular arousal and interoceptive awareness. Brain Research 1141, 178–187. 10.1016/j.brainres.2007.01.026

Sallinen, M., Kaartinen, J., Lyytinen, H., 1996. Processing of auditory stimuli during tonic and phasic periods of REM sleep as revealed by event-related brain potentials. Journal of sleep research 5, 220–228.

Salvesen, L., Capriglia, E., Dresler, M., Bernardi, G., 2024. Influencing dreams through sensory stimulation: A systematic review. Sleep Medicine Reviews 74, 101908. 10.1016/j.smrv.2024.101908

Schandry, R., Sparrer, B., Weitkunat, R., 1986. From the heart to the brain: a study of heartbeat contingent scalp potentials. Int J Neurosci 30, 261–275. 10.3109/00207458608985677

Senzai, Y., Scanziani, M., 2024. The brain simulates actions and their consequences during REM sleep. 10.1101/2024.08.13.607810

Senzai, Y., Scanziani, M., 2022. A cognitive process occurring during sleep is revealed by rapid eye movements. Science 377, 999–1004. 10.1126/science.abp8852

Seth, A.K., 2013. Interoceptive inference, emotion, and the embodied self. Trends in cognitive sciences 17, 565–573.

Silvani, A., 2008. Physiological sleep-dependent changes in arterial blood pressure: central autonomic commands and baroreflex control. Clin Exp Pharmacol Physiol 35, 987–994. 10.1111/j.1440-1681.2008.04985.x

Silvani, A., Dampney, R.A.L., 2013. Central control of cardiovascular function during sleep. American Journal of Physiology-Heart and Circulatory Physiology 305, H1683–H1692. 10.1152/ajpheart.00554.2013

Simor, P., Bogdány, T., Bódizs, R., Perakakis, P., 2021a. Cortical monitoring of cardiac activity during rapid eye movement sleep: the heartbeat evoked potential in phasic and tonic rapid-eye-movement microstates. Sleep 44, zsab100.

Simor, P., Szalárdy, O., Gombos, F., Ujma, P.P., Jordán, Z., Halász, L., Er\Hoss, L., Fabó, D., Bódizs, R., 2021b. REM Sleep Microstates in the Human Anterior Thalamus. Journal of Neuroscience 41, 5677–5686.

Simor, P., van der Wijk, G., Nobili, L., Peigneux, P., 2020. The microstructure of REM sleep: Why phasic and tonic? Sleep Medicine Reviews 52, 101305. 10.1016/j.smrv.2020.101305

Smith, C.T., 2004. Posttraining increases in REM sleep intensity implicate REM sleep in memory processing and provide a biological marker of learning potential. Learning & Memory 11, 714–719. 10.1101/lm.74904

Szalárdy, O., Simor, P., Ujma, P.P., Jordán, Z., Halász, L., Erőss, L., Fabó, D., Bódizs, R., 2024. Temporal association between sleep spindles and ripples in the human anterior and mediodorsal thalamus. Eur J of Neuroscience ejn.16240. 10.1111/ejn.16240

Takahara, M., Nittono, H., Hori, T., 2006. Effect of voluntary attention on auditory processing during REM sleep. Sleep 29, 975–982.

Takahara, M., Nittono, H., Hori, T., 2002. Comparison of the event-related potentials between tonic and phasic periods of rapid eye movement sleep. Psychiatry and clinical neurosciences 56, 257–258.

Team (2020), J., 2020. JASP (Version 0.14. 1)[Computer Software].

Tiemann, L., Schulz, E., Gross, J., Ploner, M., 2010. Gamma oscillations as a neuronal correlate of the attentional effects of pain. PAIN 150, 302–308. 10.1016/j.pain.2010.05.014

van Es, M.W.J., Schoffelen, J.-M., 2019. Stimulus-induced gamma power predicts the amplitude of the subsequent visual evoked response. Neuroimage 186, 703–712. 10.1016/j.neuroimage.2018.11.029

van der Helm, E., Yao, J., Dutt, S., Rao, V., Saletin, J.M., Walker, M.P., 2011. REM Sleep Depotentiates Amygdala Activity to Previous Emotional Experiences. Current Biology 21, 2029–2032. 10.1016/j.cub.2011.10.052

Vertes, R.P., Albo, Z., Di Prisco, G.V., 2001. Theta-rhythmically firing neurons in the anterior thalamus: implications for mnemonic functions of Papez’s circuit. Neuroscience 104, 619–625.

Wehrle, R., Kaufmann, C., Wetter, T.C., Holsboer, F., Auer, D.P., Pollmächer, T., Czisch, M., 2007. Functional microstates within human REM sleep: first evidence from fMRI of a thalamocortical network specific for phasic REM periods. European Journal of Neuroscience 25, 863–871.

Wei, Y., Van Someren, E.J., 2020. Interoception relates to sleep and sleep disorders. Current Opinion in Behavioral Sciences 33, 1–7. 10.1016/j.cobeha.2019.11.008

Zhang, Z.G., Hu, L., Hung, Y.S., Mouraux, A., Iannetti, G.D., 2012. Gamma-Band Oscillations in the Primary Somatosensory Cortex—A Direct and Obligatory Correlate of Subjective Pain Intensity. J. Neurosci. 32, 7429–7438. 10.1523/JNEUROSCI.5877-11.2012

